# An ultrasensitive spatial tissue proteomics workflow exceeding 100 proteomes per day

**DOI:** 10.1101/2025.06.02.657389

**Authors:** Melissa Klingeberg, Christoph Krisp, Sonja Fritzsche, Simon Schallenberg, Daniel Hornburg, Fabian Coscia

**Author notes:** For correspondence: Fabian Coscia,.

## Abstract

Achieving high-resolution spatial tissue proteomes requires careful balancing and integration of optimized sample processing, chromatography, and MS acquisition. Here, we present an advanced cellenONE protocol for loss-reduced tissue processing and compare all Evosep ONE Whisper Zoom gradients (20, 40, 80, and 120 samples per day), along with three common DIA acquisition schemes on a timsUltra AIP mass spectrometer. We found that tissue type was as important as gradient length and sample amount in determining proteome coverage. Moreover, the benefit of increased tissue sampling was gradient- and dynamic range-dependent. Analyzing mouse liver, a high dynamic range tissue, over tenfold more tissue sampling led to only ~30% gain in protein identification for short gradients (120 SPD and 80 SPD). However, even the lowest tested tissue amount (0.04 nL, 40,000 µm^3^) yielded 3,200 reproducibly quantified proteins for the 120 SPD method. Longer gradients (40 SPD and 20 SPD) instead significantly benefited from more tissue sampling, quantifying over 7,500 proteins from 0.5 nL of tonsil T-cell niches. Finally, we applied our workflow to a rare squamous cell carcinoma of the oral cavity, uncovering disease-associated pathways and region-specific protein level changes. Our study demonstrates that more than 100 high-quality spatial tissue proteomes can be prepared and acquired daily, laying a strong foundation for cohort-size spatial tissue proteomics in translational research.

## Introduction

Spatial proteomics (SP) has emerged as a powerful method for studying health and disease mechanisms, offering unprecedented insights into the spatial organization of proteins within cells and tissues. This rapidly evolving field has gained significant attention, with Nature Methods recognizing it as the “Method of the Year 2024” (1). Recent advances in ultrasensitive mass spectrometry have paved the way for powerful multiscale SP by combining complementary imaging and exploratory mass spectrometry (MS) approaches (2, 3). For example, we recently pioneered deep visual proteomics (4), which combines whole-slide imaging, machine-learning guided image analysis, single-cell laser microdissection (LMD) and ultrasensitive mass spectrometry. Other MS-based SP methods include nanoPOTS (5), SCPro (6), FAXP (7) or MALDI imaging guided LC-MS analysis (8). These methods rely on loss-reduced and automated sample processing for LC-MS analysis, as well as advanced MS sensitivity and throughput. To automate tissue processing following LMD, we recently developed a robotic sample processing protocol based on the cellenONE system, which enables the preparation of hundreds of tissue samples per day (9). Introducing the proteoCHIP EVO 96 for tissue proteomics following LMD, we quantified over 2,000 proteins from tissue microregions as small as 0.04 nL (40,000 µm^3^) of human tonsil tissue with a throughout of 30-40 measurements daily. Here, we expand on these efforts and derive a significantly improved protocol for more efficient and larger-scale tissue processing. We also systematically compared all Evosep ONE Whisper Zoom methods (20, 40, 80, and 120 SPD) in combination with the timsUltra AIP mass spectrometer for their suitability to deliver high-quality spatial tissue proteomes. Based on an optimized LC-MS method set tailored to ultra-low input and high throughput tissue analysis, we demonstrated that more than hundred spatially resolved proteomes can be prepared and measured daily. We validated our pipeline in a rare cancer of the oral cavity from which we profiled 170 microregions, thereby shedding light on intratumoral proteome heterogeneity. These results provide practical guidelines for the design of spatial proteomic studies and provide a solid foundation for cohort-size spatial tissue proteomics in basic and translational research.

## Results

### Streamlined workflow for high-throughput spatial tissue proteomics

To substantially increase the throughput and sensitivity of sample processing and LC-MS acquisition for low-input spatial tissue proteomics, we refined our recently developed cellenONE sample preparation workflow and assessed its utility in combination with all four Evosep ONE Whisper Zoom gradients on the latest generation timsUltra AIP mass spectrometer (**Fig. 1a**). While the Whisper Zoom LC methods (20, 40, 80, and 120 samples per day) were designed to balance sample throughput and sensitivity, the timsUltra AIP features the latest trapped ion mobility technology in conjunction with an advanced athena ion processor (AIP) utilizing programmable mass range transfers, optimized for information-rich areas of fragment ions (**Methods**). Together, this setup significantly boosts the sensitivity of deep-proteomic analysis with minimal sample input.

**Fig 1:**
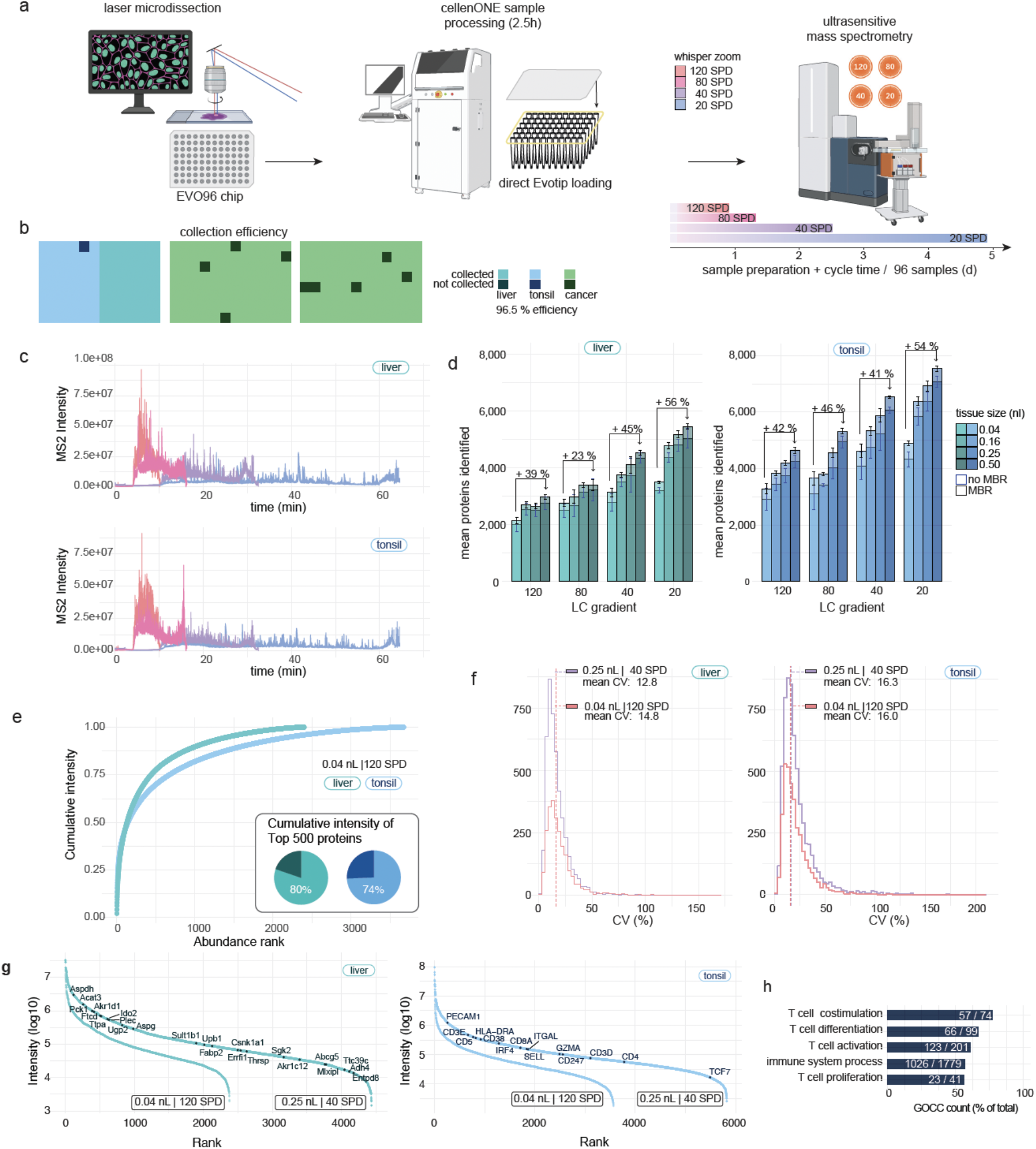
Streamlined workflow for high-throughput spatial tissue proteomics. (a) Overview of the spatial tissue proteomics workflow. (b) Collection efficiency of different 0.04 nL FFPE tissue types into the EVO 96 chip. (c) MS2 intensity of representative liver (top) and tonsil (bottom) samples (0.04 nL) measured in 120 (orange), 80 (pink), 40 (purple), and 20 (blue) samples per day. (d) Protein identification from tissue samples measured with different Whisper Zoom gradients. Volumes of 0.04 nL, 0.16 nL, 0.25 nL and 0.5 nL were laser microdissected. Averages are shown for six to eight measurements. Data were processed without (blue) and with (black) match-between-runs (MBR). (e) Cumulative protein intensities of liver and tonsil tissue proteomes (0.25 nL) ranked from the highest to the lowest abundant protein. Pie charts show the cumulative intensity of the top 500 most abundant proteins, expressed as a percentage of the total protein intensity in each sample for both tissues. (f) Coefficient of variation (CV) distributions for protein quantities across both tissue types, each represented by 0.04 nL sample in 120 SPD (orange) and 0.25 nL sample in 40 SPD (purple). The mean CV values are shown as dashed lines. (g) Dynamic range of protein abundance for liver (left) and tonsil (right) tissue, each represented by 0.04 nL sample in 120 SPD and 0.25 nL sample in 40 SPD. Hepatocyte- and T-cell-specific markers are highlighted. A minimum of two quantified values per quadruplicate measurement were required for the single-contour samples. (h) Protein identification from the immune system and T-cell specific Gene Ontology Cellular Component (GOCC).

For sample preparation, we slightly modified our previously developed protocol to make it more independent of climatic conditions because room temperature and humidity can influence sample evaporation. Therefore, we increased the rehydration volumes during the lysis and digestion steps from 500 nL to 650 nL and 150 nL to 180 nL, respectively, and supplemented the digestion buffer with 100 mM TEAB and 30% ACN. Optionally, a liquid dispenser (e.g., Formulatrix MANTIS) can be used for buffer addition to significantly speed up the protocol. These workflow refinements and the centrifugation-based Evotip loading procedure allowed us to reduce the total processing time from laser microdissection to MS acquisition-ready samples to only 2.5 h. We also refined the LMD adapter for the latest generation proteoCHIP EVO 96 (**Suppl. Fig. 1a**) and provide an updated template for 3D printing. The adapter’s close proximity to the tissue specimen inside the LMD7 microscope facilitates high tissue collection rates of > 90% for the tested tissue microregions of 0.04 nL (40,000 µm^3^, which is 8,000 µm^2^ of a 5 µm thick section) using three different tissue types (**Fig. 1b**). To perform a systematic evaluation of all four Whisper Zoom gradients for low-input tissue proteomics (**Fig. 1c**), we selected two tissue types with different dynamic ranges. Mouse liver FFPE tissue represents a more homogeneous tissue type and shows a relatively large dynamic range of protein abundance. In contrast, human tonsil tissue represents a highly structured tissue composed of spatially defined T-cell, B-cell, and squamous epithelial cell regions. Hence, this tissue type is ideal for assessing our workflow’s capacity to quantify cell type-specific proteomes. For both sample types and all four Whisper Zoom methods, we analyzed four volumes ranging from 0.04 nL to 0.5 nL of laser microdissected tissue (**Suppl. Fig. 1b-c**). The proteomic results using DIA-NN (10) are summarized in **Table 1**. As expected, longer gradients (i.e., 20 SPD and 40 SPD) resulted in higher proteome coverage for both tissue types, peaking at 7,000–8,000 proteins and 90,000-100,000 precursors from 0.5 nL of tonsil tissue (**Fig. 1d, Suppl. Fig. 1d**). However, the benefit of longer gradients was clearly dependent on the tissue type and sample amount. For example, using the 20 SPD method, ~six-fold more tissue sampling (0.04 nL versus 0.25 nL) resulted in a 48% increase in protein identification. This compared to only 24% more proteins using the 120 SPD method. For the lowest tested tissue amount (0.04 nL), we noted that the gain in throughput from short gradients (80 SPD and 120 SPD) generally outweighed the improvement in protein identification from longer gradients. In other words, a 6.6-fold longer LC gradient (68 min for 20 SPD *versus* 10.3 min for 120 SPD) led to only 37% and 46% more proteome coverage in liver and tonsil, respectively. From only 0.04 nL liver tissue, equivalent to approx. 10-15 hepatocytes (11), we quantified on average 15,545 precursors and 2,182 proteins using the Whisper Zoom 120 SPD method. This compared to 21,580 precursors and 2,904 proteins in T-cell enriched tonsil regions, which we attributed to their different dynamic ranges of protein abundance (**Fig. 1e**). Compared with our first-generation workflow (9), these data showed a three-fold increase in sample throughput at a similar proteomic depth or doubled proteome coverage (~2,000 *versus* ~4,000 proteins) at a comparable LC-MS throughput. Notably, irrespective of the tissue amount and gradient length, quantitative reproducibility was excellent for all measurements. For liver tissue, the average coefficient of variation (CV) of intra-tissue replicates was 14.8% for the smallest (0.04 nL) and 12.8 % for the largest (0.5 nL) amounts, respectively (**Fig. 1f**). For tonsil tissue, the average CVs were slightly higher at 16.3% and 16.0%, likely reflecting greater tissue heterogeneity. This trend is also evident in the global proteome correlation of around 0.96 in liver and 0.94 in tonsil, indicating high reproducibility and robustness (**Suppl. Fig. 1e**). Importantly, we quantified many known hepatocyte- and T-cell-specific markers, including known immune effector molecules (e.g., GZMA), cell type markers (CD3, CD8, and CD4), and transcription factors (e.g., TCF7 and IRF4) (**Fig. 1g**). Notably, this depth and cell-type specificity captured 1026 (58%) of all 1779 human immune system process (GOBP)-annotated proteins and over 50% of T-cell specific pathways from a single 20 SPD injection (**Fig 1h**).

**Table 1:**
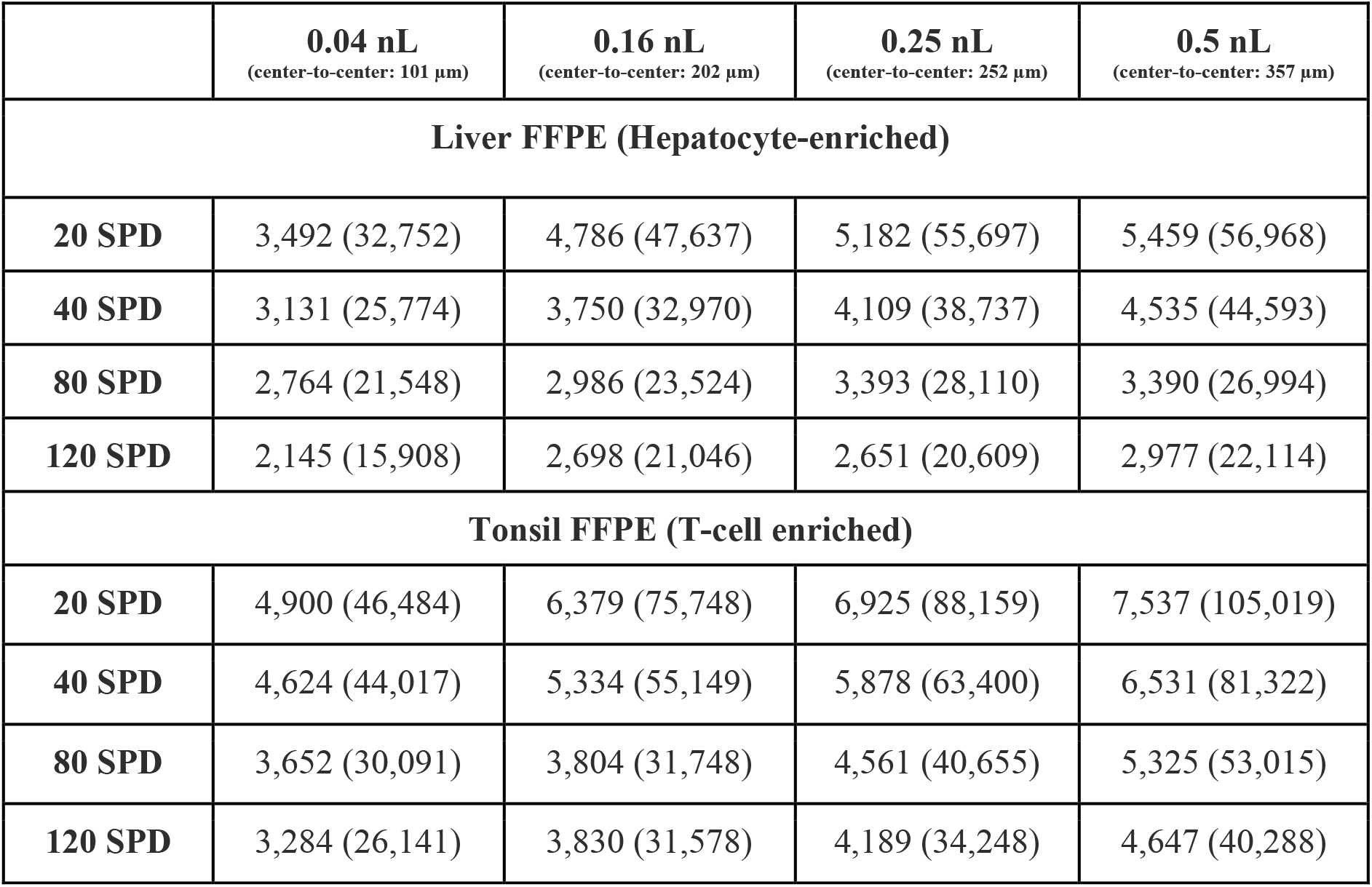
Protein and precursor identifications from four Whisper Zoom methods based on four tissue amounts and two tissue types.

| | **0.04 nL**<br>(center-to-center: 101 μm) | **0.16 nL**<br>(center-to-center: 202 μm) | **0.25 nL**<br>(center-to-center: 252 μm) | **0.5 nL**<br>(center-to-center: 357 μm) |
| --- | --- | --- | --- | --- |
| **Liver FFPE (Hepatocyte-enriched)** |  |  |  |  |
| **20 SPD** | 3,492 (32,752) | 4,786 (47,637) | 5,182 (55,697) | 5,459 (56,968) |
| **40 SPD** | 3,131 (25,774) | 3,750 (32,970) | 4,109 (38,737) | 4,535 (44,593) |
| **80 SPD** | 2,764 (21,548) | 2,986 (23,524) | 3,393 (28,110) | 3,390 (26,994) |
| **120 SPD** | 2,145 (15,908) | 2,698 (21,046) | 2,651 (20,609) | 2,977 (22,114) |
| **Tonsil FFPE (T-cell enriched)** |  |  |  |  |
| **20 SPD** | 4,900 (46,484) | 6,379 (75,748) | 6,925 (88,159) | 7,537 (105,019) |
| **40 SPD** | 4,624 (44,017) | 5,334 (55,149) | 5,878 (63,400) | 6,531 (81,322) |
| **80 SPD** | 3,652 (30,091) | 3,804 (31,748) | 4,561 (40,655) | 5,325 (53,015) |
| **120 SPD** | 3,284 (26,141) | 3,830 (31,578) | 4,189 (34,248) | 4,647 (40,288) |

### An optimized DIA method for high throughput spatial tissue proteomics

Based on our observation that the 120 SPD method provided an excellent trade-off between sample throughput and proteome coverage, we additionally evaluated six MS methods (**Fig. 2a, Table 2**) to further improve the quantitative performance of our enhanced throughput strategy. The six MS methods represented two diaPASEF-, two diagonalPASEF-, and two pyDIAid-methods with different accumulation and ramp times (75 ms and 100 ms). Similar to our gradient comparisons (**Fig. 1c-f**), we used tissue microregions of the mouse liver and human tonsil, but this time focused only on the smallest tissue regions of 8,000 µm^2^, providing a spatial resolution of ~100 µm (center-to-center distance). Our goal was to derive an optimized LC-MS method set facilitating high sample throughput, spatial resolution, and quantitative robustness without the need for excessive LMD sampling. Using our 2.5 hours cellenONE protocol and the 120 SPD setup, we completed the entire experiment including eight intra-tissue replicates for each MS method in less than one day. The experiment was conducted twice independently to increase the replicate size and validate the reproducibility and robustness of our workflow.

**Table 2:** Comparison of six DIA methods in combination with the Whisper Zoom 120 SPD method.

| no. | name | window design | accu/ramp time |
| --- | --- | --- | --- |
| 1 | diaPASEF | 6 x 3 m/z windows, 25 Da fixed | 100 ms |
| 2 |  | 8 x 3 m/z windows, 25 Da fixed | 75 ms |
| 3 | diagonalPASEF | 5 diagonal windows, 50 Da fixed | 100 ms |
| 4 |  |  | 75 ms |
| 5 | pyDIAid | 5 x 3m/z windows, variable | 100ms |
| 6 |  |  | 75ms |

**Fig 2:**
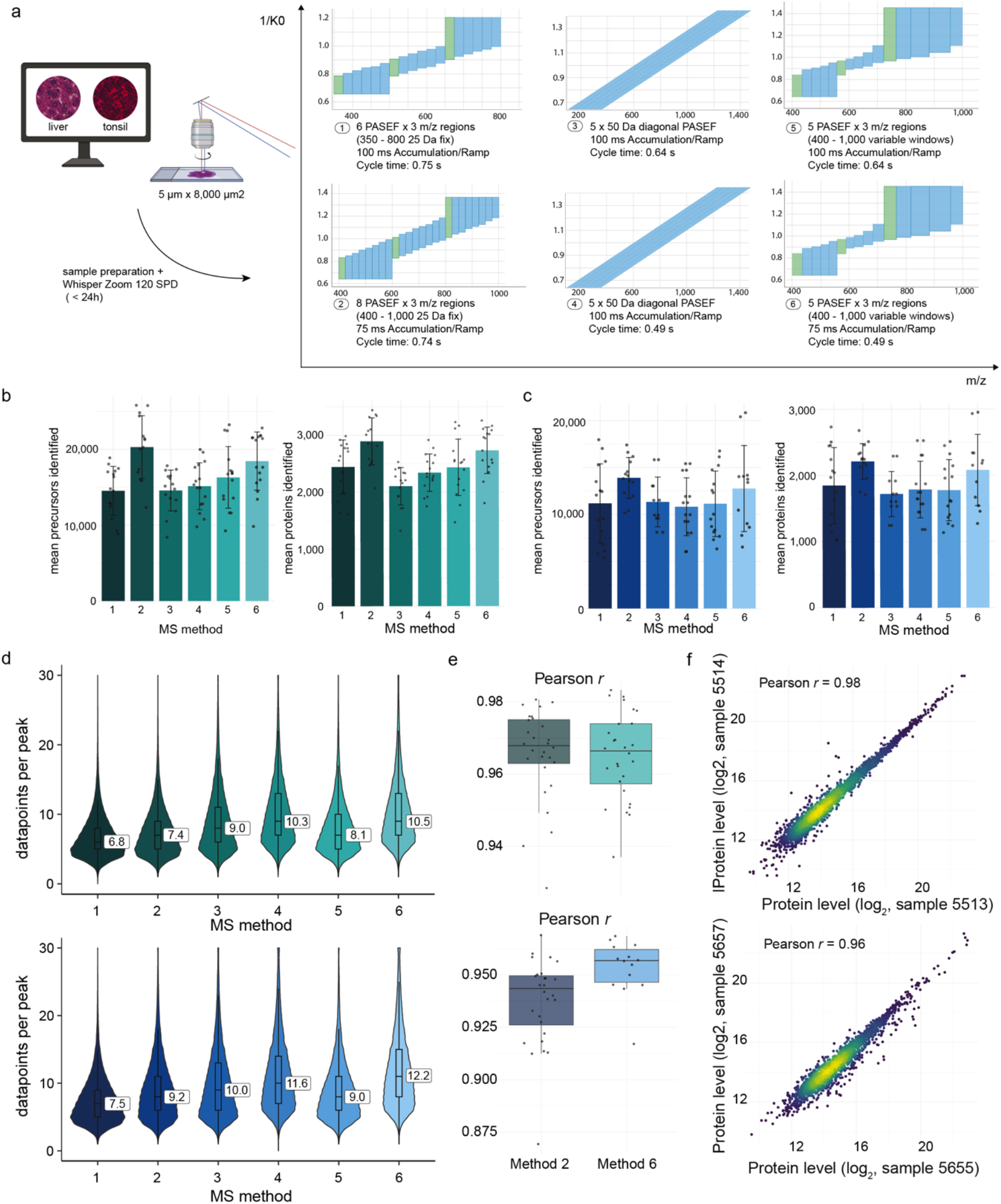
Robust and reliable protein quantification in FFPE tissue using a pyDIAid optimized method. (a) Tissue preparation strategy for benchmarking experiments using liver and tonsil FFPE tissues. Small tissue areas (0.04 nL) were laser microdissected and measured at 120 SPD, comparing six common MS methods. (b) Precursor and protein identification from liver tissue samples measured using different MS methods. Averages are shown from 16 measurements from two different experiments. (c) Precursor and protein identifications from T-cell-rich tonsil tissue samples measured using different MS methods. Averages are shown from 16 measurements from two individual experiments. (d) Violin plots display the distribution of data points per peak for each MS method in the liver (top) and tonsil (bottom) tissues. The overlaid boxplots show the median (central line), interquartile range (box), and whiskers extending to 1.5 × the IQR. (e) Boxplots showing proteome correlations (Pearson’s r) for methods 2 and 6 of one experiment. (f) Proteome correlation (Pearson’s r) between two representative samples.

Overall, we found that the chosen MS method had a significant impact on the number of identified proteins in both tissues, with an average increase of 37% and 29% comparing the best and the worst method in the liver and tonsil tissue, respectively (**Fig. 2b-c**).

Except for diagonalPASEF, a shorter accumulation time resulted in significantly higher protein identification, with an average of 2,700 proteins in the liver and 2,200 proteins in the tonsil tissue. Both the vendor standard diaPASEF 3 × 8 (method 2) and the pyDIAid optimized method with 75 ms accumulation and ramp time (method 6) resulted in the highest identification rates. As an additional metric, we evaluated data points per peak to assess the quantitative performance of the six methods. The faster diagonalPASEF (method 4) and pyDIAid optimized methods (method 6) featured the highest average number of 10–12 data points per peak (**Fig. 2d**). Considering proteome coverage and data points per peak, we concluded that method 6, the pyDIAid-optimized window setup with 75 ms accumulation and ramp time, offers the best overall performance. However, the standard 3 × 8 diaPASEF method (75 ms accumulation and ramp time) performed second best, also offering an excellent choice. This was also reflected in high global proteome correlations of 0.97 and 0.96 in liver and 0.94 and 0.95 in tonsil tissue replicates (**Fig. 2e-f**). These two methods support robust and reliable protein identification, thus making them well suited for high-throughput applications to uncover spatial tissue proteome heterogeneity.

### Tissue-wide proteome profiling of a rare squamous cell carcinoma of the oral cavity

Based on our newly developed high-throughput spatial tissue proteomics workflow, we set out to study a rare and largely understudied squamous cell carcinoma (SCC) of the oral cavity. Following IF (panCK, COL1A1, CD3 and CD20) and H&E staining, we performed tissue-wide profiling of small ROIs to analyze SCC intratumoral proteome heterogeneity. We systematically sampled 192 microregions (8,000 µm^2^) of the stromal and cancer compartments guided by IF and H&E, distributed over a whole tissue section of two centimeters in diameter (**Fig 3a**). To assess the robustness of our sample preparation protocol, the samples were processed in two separate cellenONE experiments. Sample preparation and LC-MS analysis took less than two days and resulted in a total of 170 high-quality proteomic measurements after filtering (**Fig 3b-c**). However, using SCC tissue, we noted that gravity-based tissue collection was less efficient than seen in our liver and tonsil experiments, resulting in a fraction of tissue contours that did not end up in the well bottom, resulting in low-quality proteomes. We therefore reintroduced an organic solvent rinsing step prior to tissue lysis to ensure optimal proteomic results, reducing sample loss from 19% to only 4% (**Fig 3b, Methods**). As the collection efficiency is highly tissue-and cell type-specific, we generally recommend such an organic solvent wash for enhanced consistency.

**Fig 3:**
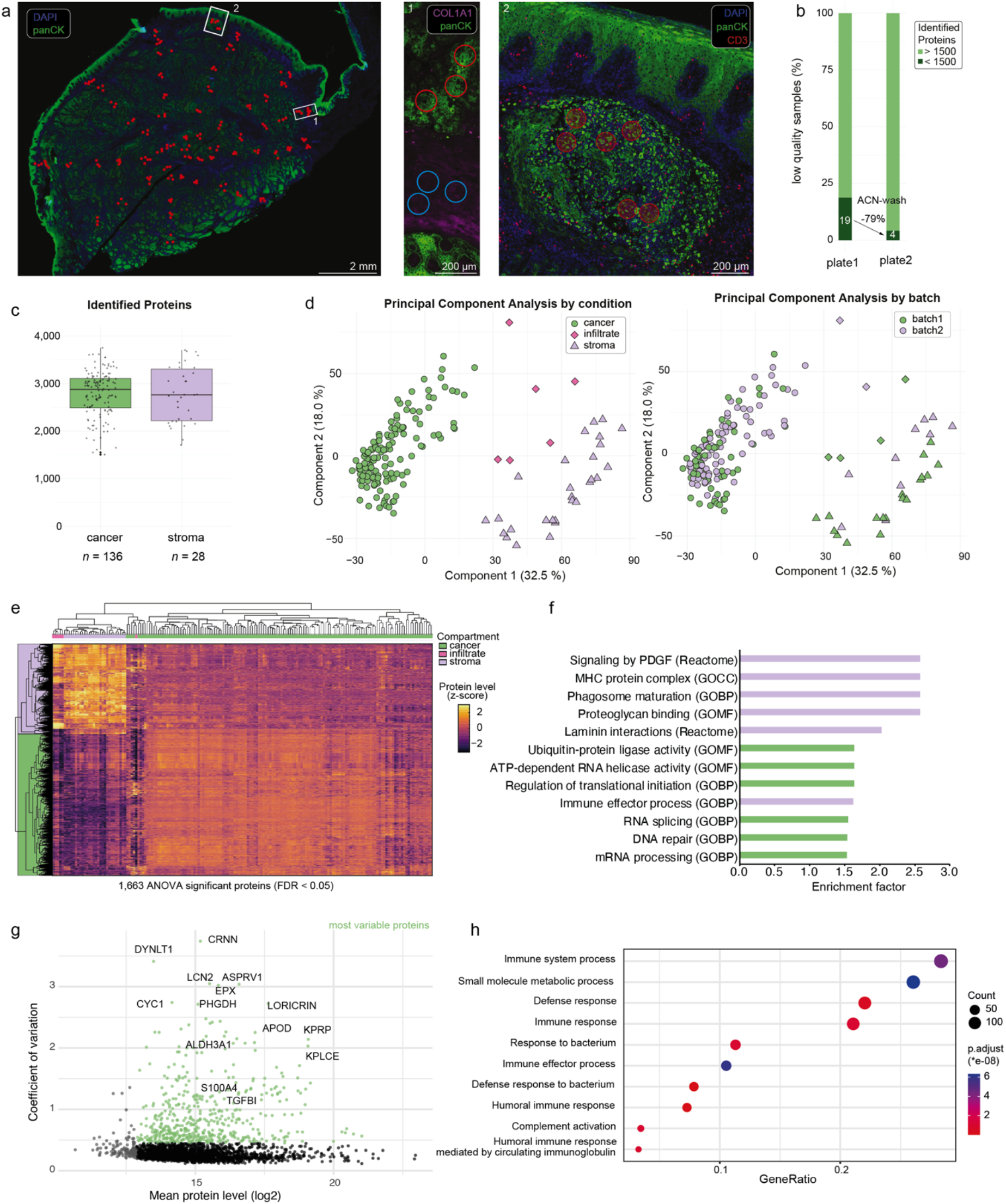
Tissue wide profiling of a head and neck squamous cell carcinoma. (a) left: Immunofluorescent whole-slide image of a 5 µm-thick squamous cell carcinoma tissue section. Red circles show regions of tissue collection; middle: Magnification of exemplary stromal and tumor area with annotations used for laser microdissection and proteomics profiling; right: T-cell infiltrated tumor niche. (b) Number of high-quality proteomes from two experiments with and without Acetonitrile wash. (c) Protein identifications from cancer and stroma tissue samples. (d) Principal-component analysis (PCA) of 170 samples after 70% valid value filtering and imputation by condition (left) and batch (right). (e) Unsupervised hierarchical clustering of ANOVA significant proteins (p < 0.05) across all samples (f) Gene Ontology (GO) enrichment analysis on ANOVA significant protein in (e). (g) Coefficient of variation plot of cancer proteome. Most variable proteins (top quintile) are marked in green. (h) Pathway enrichment analysis of variable proteins from (g).

Using the 120 SPD method, we identified on average 2,812 and 2,719 proteins in cancer and stromal regions, respectively, and 5,453 proteins in total (**Fig 3c**). Proteomes showed high compartment specificity and clustered by region-specific cellular composition. Notably, principal component analysis (PCA) revealed no detectable batch effect, underlining the robustness of our pipeline (**Fig. 3d**). Unsupervised hierarchical clustering of the 1,663 ANOVA significant proteins between sample groups (tumor, stroma, and immune infiltrate) revealed distinct expression patterns across tissue compartments (**Fig. 3e**). Interestingly, we identified a small niche of T-cell infiltrated cancer cells surrounded by stromal cells, suggesting that these regions exhibited a distinct phenotype with unique tumor-immune and tumor-stromal interactions (**Fig 3a**). Their proteomes were clearly distinct from the main tumor area and grouped closely with the stromal samples, indicative of a more mesenchymal phenotype (**Fig. 3e**). To gain further insights into the upregulated biological processes in this tumor disease, we performed pathway enrichment analysis for the two main clusters (**Fig. 3f**). Stromal proteomes showed high levels of biological processes such as signalling of PDGF, MHC protein complex and laminin interactions. In the tumor compartment, most enriched pathways included known hallmark features of cancer biology, including protein translation, ubiquitin-protein ligase activity, RNA splicing and DNA repair. To assess intra-tumoral proteome heterogeneity, we calculated the protein-level variability (coefficient of variation, CV) across more than 140 samples that were distributed over the entire tissue section (**Fig. 3g**). We identified dozens of proteins with high protein level variability across the different cancer regions, including CYC1 and ALDH3A1, suggesting spatial differences in metabolic activity and oxidative stress adaptation. Notably, we identified cornulin (CRNN), also known as squamous heat shock protein 53, as the most variable protein in our dataset. Cornulin was identified to have cancer-promoting effects in SCC (12) and may reflect spatial differences in cell differentiation. The high variability of EPX, a granulocyte marker, indicated pronounced differences in the tumor-immune microenvironment in this tumor. Affirmatively, pathway enrichment analysis of the 20% most variable proteins identified immune system and innate immunity related processes as most significantly enriched together with metabolic changes (**Fig. 3h**).

In summary, our spatially resolved proteomic data uncovered disease-associated pathways and nominated proteins associated with high region-specific expression and intratumoral heterogeneity.

## Discussion

The field of spatial proteomics is rapidly advancing and driven by innovative methodologies that integrate imaging-based cell phenotyping, laser microdissection, and exploratory mass spectrometry-based proteomics (2). To facilitate the widespread adoption of these approaches and to make them amenable to cohort-scale proteome profiling, improvements in sensitivity, throughput, and spatial resolution are essential. This not only necessitates robust and automated sample preparation workflows capable of efficiently processing hundreds of samples, but also balanced LC-MS strategies that offer high sensitivity and throughput. In this study, we refined our previously developed cellenONE tissue protocol and systematically evaluated all Evosep ONE Whisper Zoom gradients (20–120 SPD) on the timsUltra AIP mass spectrometer based on liver and tonsil FFPE tissues. We demonstrated that short gradients (e.g., 120 SPD) provide an excellent compromise between throughput and proteomic depth for ultra-low tissue amounts, enabling the preparation and acquisition of more than 100 spatially resolved proteome measurements daily. We quantified over 3,000 proteins from as little as 0.04 nL of FFPE tissue with excellent quantitative reproducibility. Generally, the benefit of longer gradients (e.g., 20 SPD) was most apparent when combined with increased tissue sampling. For instance, following whole-slide immunofluorescence imaging, we profiled over 7,500 protein groups from 0.5 nL of tonsil tissue per single 20 SPD injection. Notably, this depth was sufficient to capture more than half of all immune system process (GOBP)-related proteins annotated in the genome. In addition, we observed that tissue type is an equally important factor defining proteome coverage as gradient length and tissue input amount. In contrast to our tonsil results, using liver tissue, we observed that protein identification saturated around 5,000–6,000 proteins, despite increased tissue sampling by LMD. These findings have considerable implications for the future design of spatial proteomic studies. While high dynamic range tissues such as the liver on the one hand offer an exciting entry point for applications with excellent spatial resolution, such as single-cell applications (13, 14), proteome-wide measurements (e.g., 7,000 - 10,000 proteins) are comparably challenging to achieve from single-shot analyses and may require additional fractionation methods, such as gas-phase fractionation techniques (15) to reduce sample complexity. This highlights the importance of balancing the excised tissue area and the number of dissected contours based on the underlying tissue properties. These results raise some fundamental questions regarding spatial proteomics study design. For example, how much tissue needs to be sampled to address the biological question of interest the best? Key parameters to consider are, (1) dynamic range distribution of protein abundance (i.e., how “expensive” are the next 500 proteins), (2) spatial gradients within tissues (how much biological granularity is needed, choosing between single-cell LMD approaches or multicellular regions/niches), culminating in the question (3) to balance the number of tissue sections, contour size and gradient.

We also compared three common DIA schemes and derived an optimized pyDIAid method that achieved superior proteome coverage and quantitative performance for limited tissue amounts. Our analysis comparing different DIA methods also highlighted the importance of optimizing methods specifically for tissue samples, rather than relying solely on cell culture evaluations (e.g., HeLa peptide dilutions). Together with our Whisper Zoom gradient tests, we derived an optimized method set for 120 high-quality spatial proteomics per day. We demonstrated the feasibility of our optimized end-to-end workflow for large-scale and tissue-wide proteome profiling of a rare squamous cell carcinoma of the oral cavity. Guided by IF microscopy, we profiled 170 microregions and shed light on intratumoral proteome heterogeneity in this largely under-studied cancer. Our results revealed an upregulation of proteins related to translation, the ubiquitin system, RNA splicing, and DNA repair, and identified dozens of proteins showing high intratumoral heterogeneity, thus providing a valuable resource for SCC future studies.

In summary, this study presents an optimized and scalable workflow for large-scale spatial tissue proteomics, laying a strong foundation for spatial discovery proteomics in basic and translational research.

## Experimental Procedures

### Mouse Liver Tissue Collection

All animal experiments were performed according to the United Kingdom Coordinated Committee on Cancer Research (UKCCR) guidelines with approval by the local governmental authorities (Landesamt für Gesundheit und Soziales Berlin, approval number G0004/14). C57BL/6 mice from Jackson Laboratory were used and housed in individually ventilated cages in a specific pathogen-free mouse-facility at the Max-Delbrück-Center for Molecular Medicine (Berlin, Germany). To perform liver excision, mice were anesthetized and then euthanized, after which their livers were extracted, washed twice with ice-cold PBS, and placed in a 4% formaldehyde solution for fixation, which lasted between 24 and 48 hours. Subsequently, the livers were embedded in paraffin for further histological examination.

### Human Tissue Collection

The experiments for optimizing the sample LCMS methods were performed using formalin-fixed, paraffin-embedded (FFPE) human tissue: one specimen of non-neoplastic palatine tonsil and one of squamous cell carcinoma (SCC) of the maxilla. Both samples were obtained from the archive of the Institute of Pathology at the Charité – Universitätsmedizin Berlin, Campus Benjamin Franklin, Berlin, Germany. The tonsil specimen was resected from a 26-year-old male patient with recurrent tonsillitis. After fixation in 10% neutral buffered formalin, the specimen was weighed, measured, macroscopically examined, and sliced into 5-mm sections. Two representative sections were paraffin-embedded. From the resulting tissue block, 2-mm-thick sections were cut and stained with hematoxylin and eosin (H&E). Histological evaluation revealed follicular hyperplasia of the lymphoid tissue and stromal sclerosis. The SCC specimen was derived from a 53-year-old female patient. The maxillary resection specimen was fixed, measured, and inked to mark resection margins before slicing into 5-mm sections. On sectioning, a 40 mm, white-gray, firm tumor was identified with a 2 mm distance to the anterior soft tissue margin. Representative tumor areas, including regions close to the margins, were paraffin-embedded. Sections were cut and stained with H&E. Histological and immunohistochemical analysis revealed a p16-negative, moderately differentiated keratinizing SCC with bone infiltration, focal perineural invasion, and no angioinvasion, epithelial dysplasia, or lymphovascular invasion. PD-L1 expression was quantified as follows: TC score 15%, IC score 12%, CPS 27. Final tumor staging was pT4a pN2c (2/46) L0 V0 Pn1 G2 R0.

All tissue blocks were then stored at room temperature at the archive of the Institute of Pathology at Charité University Hospital, Campus Benjamin Franklin. The study was performed according to the ethical principles for medical research of the Declaration of Helsinki and approval was obtained from the Ethics Committee of the Charité University Medical Department in Berlin (EA1/222/21).

### Tissue sectioning, Immunofluorescent staining and H&E staining

The FFPE blocks were sectioned at 5 μm thickness on PPS frame slides (Leica, 11600294) and left in the oven overnight at 37 °C. Slides were heated at 60 °C for 30 min to improve tissue adhesion and deparaffinized. Mouse liver samples were stained with H&E. Human Tonsil and HNSCC samples were stained by immunofluorescence. To improve antibody binding, the samples underwent a process of heat-induced epitope retrieval. In summary, they were heated to 95 °C for 30 minutes and then allowed to cool at room temperature for 30 minutes. For tonsil tissue, three conjugated antibodies targeting CD20 (dilution 1:50, Thermo Fisher Scientific, 53-0202-80, Alexa Fluor 488), CD3 (dilution 1:100, NovusBio, NBP2-52710AF647, Alexa Fluor 647), pan-cytokeratin (dilution 1:100, Thermo Fisher Scientific, 41-9003-82, eFluor 570) were used. For the carcinoma tissue, four conjugated antibodies targeting pan-cytokeratin, CD20, CD3 and COL1A1 (dilution 1:50, NovusBio, NB600-408AF750, Alexa Fluor 750) were used. Overnight tissue staining was conducted at 4 °C within a humidity chamber. Antibodies were diluted in 3% BSA (in 1x PBS, Serva, Cat. No. 11948.01). Following antibody incubation, tissues underwent four washes with PBS, and nuclear staining was achieved with a 10-minute Hoechst treatment (1:1000 in PBS, Thermo Fisher Scientific, 62249). To improve well inspection for laser microdissection, tissues were H&E stained.

### Design of Leica LMD7 Collection Plate Adapters

We designed a new adapter for the new version of the proteoCHIP EVO 96. The adapter was crafted from transparent acrylic glass (PMMA-gs) using a CNC milling machine. It can also be manufactured with standard 3D printers using the supplied files (.stl format). The design and measurements are depicted in Supplemental Figs. S1, and the 3D printer files are available in the Supplementary Materials.

### Whole-Slide Imaging and LMD

Human tissue sections labeled with immunofluorescence were imaged using a Zeiss Axioscan 7 system, which features wide-field optics, a Plan-A photochromat 10x/0.45 M27 objective, and a quadruple-band filter set for Alexa fluorescent dyes. These images were then imported into QuPath (version 0.5.1) for further annotation. A trained pathologist collaborated on annotating the various regions of interest. The annotations, along with three reference points for contour alignment, were exported in geojson format and converted to the necessary .xml format for LMD. The contours were mapped to specific target locations on the proteoCHIP EVO 96. The code for processing these shapes is accessible at github.com/CosciaLab/Qupath_to_LMD, utilizing geopandas and the py-lmd package. LMD was performed using the Leica LMD7 system with Leica Laser Microdissection software V 8.3.0.08259. A customized plate layout was defined by using the universal holder function in the LMD software. Depending on the contour size, tissue was cut with a 10x or 20x dry objective in brightfield mode.

### Sample Preparation Using the cellenONE System

To concentrate tissue samples at the bottom of the proteoChip EVO 96, 10 μL of acetonitrile can be introduced into each well post-collection and allowed to air dry at room temperature for 10 minutes. It is advisable to conduct another well inspection before preparing proteomics samples to ensure efficient collection. For the cellenONE, water that is purified and filtered (>18 MΩ, <3 ppb TOC at 25 °C) was utilized. Two microliters of lysis buffer (comprising 0.1% n-dodecyl β-D-maltoside (DDM), 0.1 M tetraethylammonium bromide at pH 8.5 (TEAB), 5 mM tris(2-carboxyethyl)phosphine (TCEP), and 20 mM chloroacetamide (CAA)) was dispensed into each well using the MANTIS Liquid Dispenser (Formulatrix, V3.3 ACC RFID, software version 4.7.5) with the high-volume diaphragm chips (Formulatrix, cat.no. 233128). For the lysis process, the sample was heated within the cellenONE, which was set to operate at 65 °C and 85% humidity. Continuous rehydration (650 nL/cycle, 1000 Hz) was enabled to prevent the lysis buffer from evaporating from the wells. Note, the volume dispensed per cycle may need adjustment based on local temperature and humidity to prevent complete evaporation. After a 60-minute incubation, the temperature was lowered to 20 °C, and rehydration continued until the temperature reached 25 °C. At 25 °C, the process was halted, and one microliter of enzyme mix (10 ng/µL lysC & trypsin (Promega, Cat. V5072), 0.1 M TEAB, 30% ACN) was dispensed using the MANTIS Liquid Dispenser. The reaction mixture was incubated within the cellenONE for one hour at 37°C with 85% humidity, maintaining continuous rehydration (200 nL/cycle, 500 Hz).

### Peptide Clean-Up with C-18 tips

After digestion, peptide clean-up was performed using Evotips (EV2013, Evotip Pure, Evosep) as recommended by the manufacturer. Shortly, tips were rinsed with 20 μL of buffer B (99.9% ACN, 0.1% FA) and centrifuged at 800 rcf for 1 min. For equilibration, 20 μL of buffer A (99.9% water, 0.1% FA) was added to each tip, activated in isopropanol for 10 s, and centrifuged again at 800 rcf for 1 min. 17 µL of Buffer A was added and the proteoCHIP EVO 96 was flipped directly onto the activated Evotips. The samples were transferred by centrifugation at 800 rcf for 1 min. The wells of the Evo96 Chip were washed with 3 µL Buffer A and the sample transfer step repeated. To each tip, 100 μL of buffer A was added, followed by centrifugation at 800 rcf for 10 seconds to ensure the liquid reached the membrane. The tips were then placed in a tray with a holder containing buffer A, ensuring they remained submerged and did not dry out.

### LC–MS Analysis

LC-MS analysis was performed on Evosep ONE (Evosep Biosystems) LC system connected to a trapped ion mobility spectrometry quadrupole time-of-flight mass spectrometer (timsUltra AIP, Bruker Daltonics). The Evosep ONE was operated in the Whisper Zoom configuration and was used with Aurora Rapid 75 (5 cm x 75 µm x 1.7 µm, IonOpticks) and Aurora Elite (15 cm x 75 µm x 1.7 µm, IonOpticks) both kept at 50°C. Buffer A consists of 0.1% formic acid in LC-MS grade water and buffer B is 0.1% formic acid in acetonitrile.

Peptides were separated with Whisper Zoom 20 SPD, 40 SPD, 80 SPD and 120 SPD method. Eluted peptides were detected in dia-PASEF and diagonal-PASEF using a timsUltra AIP mass spectrometer operated in positive ion mode equipped with a captive spray Ultra 2 ion source. Capillary voltage was set to 1300 V for Whisper Zoom 120 SPD, and 80 SPD and to 1500 V for Whisper 40 SPD, and 20 SPD with a capillary temperature of 200 °C. dia-PASEF methods used were 1) a 3 m/z range, 6 MS/MS PASEF cycles with fixed 25 Da windows from 350 – 800 m/z in an ion mobility range between 0.64 – 1.25 1/k0 with 100 ms accumulation and ramping time at a cycle time of 0.75 s, 2) a 3 m/z range, 8 MS/MS PASEF cycles with fixed 25 Da windows from 400 – 1000 m/z in an ion mobility range between 0.64 – 1.45 1/k0 with 75 ms accumulation and ramping time at a cycle time of 0.74 s; 3 & 4) a 3 m/z range, 5 MS/MS PASEF cycles with variable windows optimized based on ion density distribution using pyDIAid (16) from 400– 1000 m/z in an ion mobility range between 0.64 – 1.45 1/k0 with 100 ms or 75 ms accumulation and ramping time at a cycle time of 0.64 s and 0.49 s, respectively; 5 & 6 a diagonal-PASEF method of 5 non-overlapping MS/MS PASEF cycles with slice widths of 50 Da in an ion mobility range between 0.64 – 1.45 1/k0 with 100 ms or 75 ms accumulation and ramping time at a cycle time of 0.64 s and 0.49 s, respectively.

### Raw File Processing

DIA-NN (10) Academic (version 2.0) was used for raw file analysis. DIA-NN in silico predicted libraries were generated by providing the human FASTA file and frequently found contaminants (17) (UP000005640_9606, downloaded on March 6th 2025, respectively). For mouse liver analysis, we used a tissue-specific library as described earlier (14). The refined murine liver library consisted of 68,006 precursors, 61,554 elution groups, and 8225 protein groups. DIA-NN was operated in the default mode with minor adjustments. Briefly, precursor false discovery rate (FDR) was 1%, precursor charge state 2 to 4, precursor m/z range to 300 to 1200, MS1 and MS2 accuracies 15.0 ppm, scan windows 0 (assignment by DIA-NN), MBR and protein inference were enabled. Oxidation (M) and acetyl (Protein-N-term) were included as variable modifications and carbamidomethyl (C) as fixed modification with maximum two allowed modifications per peptide. For data acquired with diagonalPASEF, the additional option –tims-scan was provided. We used the report.stat and report.pg matrix output file of DIA-NN for further data analysis with a global protein q-value threshold of 1%.

### Proteomics Data Analysis

Proteomics data analysis was performed with Perseus (18) (versions 1.6.50 and 2.1.2.0) and within the R environment (https://www.r-project.org/, version 4.4.3) with the following packages: ggplot2 (v3.5.1), dplyr (v1.1.4), tidyr (v1.3.1), stringr (v1.5.1), data.table (v1.17.0). CVs were calculated in linear space.

### Experimental Design and Statistical Rationale

For each proteomic comparison, a minimum of three replicates per sample group was employed. Replicates were defined as adjacent regions isolated from the same tissue section, exhibiting a similar cell type composition as determined by microscopic imaging. In the experiments involving murine liver and human tonsil depicted in Figure 1, six replicates were collected for each condition, defined by the combination of tissue size and LC gradient. For the MS method comparison (Fig. 2), eight replicates per tissue and method were collected, and the experiment was conducted twice, resulting in a total of 16 replicates. Coefficients of variation (CVs) were calculated in linear space. For the multi-group comparison in Figure 3, which examines tumor, stroma, and infiltration zone samples, ANOVA was employed with a permutation-based false discovery rate (FDR) of 5%. Detailed methodologies for the statistical tests applied are provided in the corresponding figure legends.

## Supporting information

EVO96-LMD7-Adapter

## Author contributions

Conceptualization: M.K., C.K., and F.C.; Methodology: M.K., C.K., S.F., S.S.; Experiments: M.K., C.K., S.F., S.S.; Data curation: M.K., C.K.; Data analysis: M.K., C.K., and F.C.; Figures: M.K.; Supervision: D.H and F.C.; Funding acquisition: F.C.; Writing the original draft: M.K and F.C. All authors reviewed and edited the manuscript.

## Acknowledgments

We would like to thank our colleagues at the Max-Delbrück-Center (MDC) for their support and fruitful discussions. In particular, we thank Steffen Tornow from the TFM/scientific workshop for his support to design the LMD adapters. Ulrike Stein, we thank for her support to perform the mouse liver experiments. Furthermore, we acknowledge the MDC technology platform ‘Proteomics’ (AG Mertins) for their great support. M.K, S.F., and F.C. acknowledge funding support by the Federal Ministry of Education and Research (BMBF), as part of the National Research Initiatives for Mass Spectrometry in Systems Medicine, under grant agreement No. 161L0222. This project received funding from the European Research Council (ERC) under the European Union’s Horizon 2020 research and innovation program (grant agreement No. 101115681) and support by the ERC (ERC starting grant). This research was supported by the Initiative and Networking Fund of the Helmholtz Association within the framework of the Transfer Campaign (Helmholtz Co-Creation Projects).

## Data availability

The mass spectrometry proteomics data have been deposited to the ProteomeXchange Consortium (http://proteomecentral.proteomexchange.org) via the PRIDE partner (19) and are available upon manuscript acceptance.

---

**Supplementary Fig. 1, related to Fig. 1:**
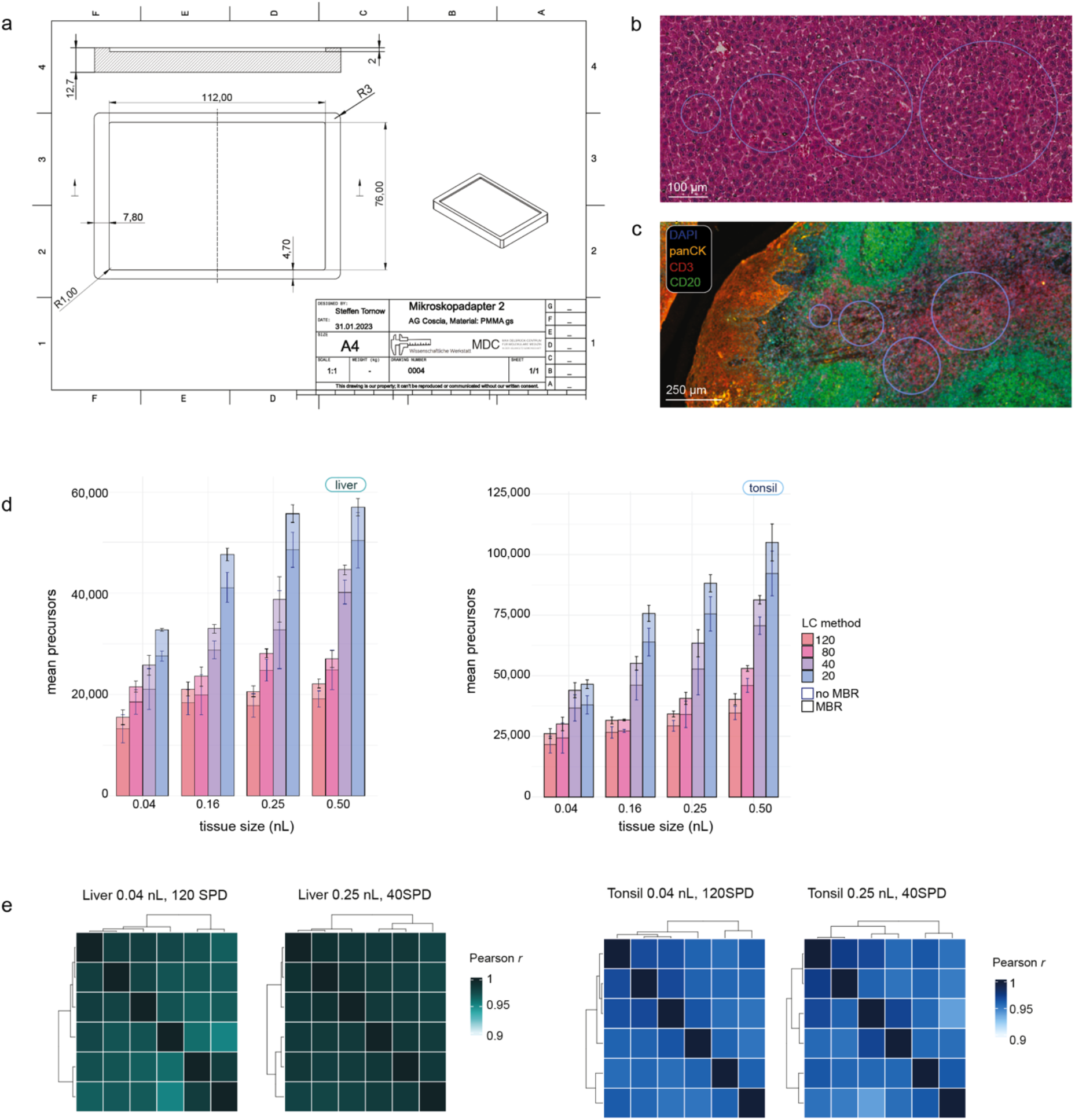
(a) Technical drawing of LMD7 adapter compatible with Evo 96 chip. (b) H&E staining of 5 µm murine liver tissue section, illustrating four isolated area sizes: 8,000 µm^2^, 32,000 µm^2^, 50,000 µm^2^, 100,000 µm^2^. (c) Immunofluorescent staining of 5 µm human tonsil tissue section, showing panCK (orange), CD3 (red), CD20 (green) and DAPI (blue). (d) Identified precursors from different sized liver (left) and tonsil (right) tissue samples measured with different Whisper Zoom methods. Mean values from six to eight replicates are shown, with (blue) and without (black) MBR. (e) Heatmap of Proteome correlation (Pearson’s r) between replicates of the same size measured with identical LC gradients for liver (left) and tonsil (right).

